# The dissemination of Rift Valley fever virus to the eye and sensory neurons of zebrafish larvae is *stat1* dependent

**DOI:** 10.1101/2022.08.10.503461

**Authors:** S. ter Horst, A. Siekierska, AS. De Meulemeester, A. Cuvry, L. Cools, J. Neyts, P. de Witte, J. Rocha-Pereira

**Affiliations:** Laboratory for Virology and Chemotherapy, Rega Institute for Medical Research, Department of Microbiology, Immunology and Transplantation, KU Leuven, Leuven, Belgium; Laboratory for Molecular Biodiscovery, Department of Pharmaceutical and Pharmacological Sciences, KU Leuven, Leuven, Belgium

**Keywords:** Bunyavirus, Rift Valley fever virus, innate immunity, zebrafish, pathogenesis

## Abstract

The Rift Valley fever virus (RVFV) is listed by the WHO as priority disease and causes haemorrhagic fever, encephalitis, and permanent blindness. To study RVFV pathogenesis and identify small-molecule antivirals, we established a novel *in vivo* model using zebrafish larvae. Pericardial injection of RVFV resulted in ^~^4 log_10_ viral RNA copies/larva, which was inhibited by antiviral 2′-fluoro-2′-deoxycytidine. The optical transparency of the larvae allowed detection of RVFV_eGFP_ in the liver and sensory nervous system, including the optic tectum and retina, but not the brain or spinal cord. Thus, RVFV-induced blindness likely occurs due to direct damage to the eye and peripheral neurons, rather than the brain. Treatment with JAK-inhibitor ruxolitinib, as well as knockout of *stat1a* but not *stat1b*, enhanced RVFV replication to ^~^6 log_10_ viral RNA copies/larva and ultra-bright livers, although without dissemination to sensory neurons or the eye, hereby confirming the critical role of *stat1* in RVFV pathogenesis.

## Introduction

Rift Valley fever virus (RVFV) (order: *Bunyavirales*, family: *Phenuiviridae*, genus: *Phlebovirus*) is a mosquito-borne pathogen able to cause acute viral hemorrhagic disease. It is included in the WHO list of priority diseases due to its potential to cause a public health emergency and the absence of efficacious vaccines and treatments. (1) The virus, with its segmented negative single-stranded RNA genome (ssRNA), can infect both ruminants and humans, and is widely endemic throughout the African continent and the Arabian Peninsula where it has already caused substantial outbreaks. Humans become infected through mosquito bites or via contact with body fluids or tissue of infected animals. Most humans infected with RVFV present symptoms such as self-limiting mild malaise, muscle pain, and fever. However, about ^~^10% of individuals develops severe symptoms such as haemorrhagic fever, encephalitis or ocular disease. (2) Case fatality in people infected with RVFV is overall ^~^1% but goes up to 50% for those who develop haemorrhagic fever. Ocular complications such as reduced vision, photophobia and retroorbital pain are frequent and may take months to resolve or result in permanent blindness. (3) While retinal haemorrhage has been observed in some patients, it is unclear whether this is the actual cause of the patients loss of sight. (4) This type of severe clinical outcome stresses the need to develop specific antiviral treatment and vaccines against RVFV infection.

A detailed understanding of viral dissemination, pathogenesis, and host immune response to a RVFV infection is necessary for the design and development of effective countermeasures. The remaining knowledge gaps include understanding how RVFV disseminates past the biting site and which cell types are involved, the nuances in pathogenesis between the multiple infected species, and which host factors limit or enable infection exacerbation. (5) Small animal models play a pivotal role in unravelling these details. The virus replicates in several species, including rodents, ruminants, and non-human primates, hence these were used in a variety of RVFV-related studies. In rodents, RVFV infection is typically lethal due to the development of hepatic necrosis, haemorrhagic disease, or encephalitis. (6–10) While they come with major practical disadvantages and ethical concerns, non-human primates models best match the severe disease phenotypes of human RVFV infection. (11)

Over the years, zebrafish have gained ground in the field of infectious diseases and particularly of human viral diseases due to its attractive features such as small size and vertebrate nature. On top of being better compliant with the 3 R’s (replace, reduce, and refine) rule, the model offers numerous advantages. As their larvae are optically transparent, they are ideally suited for live imaging and allow follow-up of virus dissemination throughout the different organs and systems. Moreover, zebrafish are easily genetically manipulated and have rapid development thus resulting in a relatively fast generation of transgenic lines bearing gene knock-ins, knockouts, and/or expressing fluorophores under tissue-specific promotors. Zebrafish models to study human viral infections have been established for human herpes simplex 1 virus, influenza virus, chikungunya virus, and sindbis virus. (12–15) More recently, our team developed a zebrafish larvae model for human norovirus infections, which constituted a breakthrough given the decades-long difficulties to cultivate this important enteric pathogen. (16) Additionally, attempts have been made to infect zebrafish larvae with SARS-CoV-2, but only limited viral replication was observed in a subset of animals when infected in through the swim bladder. (17) However, it was suggested that transgenic zebrafish stably expressing human ACE2 and/or transmembrane serine protease (TMPRSS2) have the potential to effectively host SARS-CoV-2 replication

An important feature of zebrafish larvae is that they lack a mature adaptive immune system in the first 4-6 weeks post fertilization. (18) However, the innate immune system is in place from the earliest days post fertilization (dpf), with functional macrophages and neutrophils present from 30 hours post fertilization, hence providing the unique possibility to dissect in detail the virus-related innate immune response separately. (19) To that end, transgenic lines bearing fluorescent neutrophils and macrophages are readily available, and the main functions and signalling pathways are conserved with the innate immune system of humans. (20) Among those, the interferon (IFN) system is a major driving force, since the ability of an individual mammalian cell to respond to a viral infection is based on the signalling through type I IFNs. (21) In zebrafish four types of type I *ifn* are distinguished, *ifnφ1-4*. Many of the *ifn*-stimulated genes (ISGs) produce functional and structural orthologues to human ISGs. (22) As in humans, zebrafish innate immunity against ssRNA viruses is initiated upon detection of viral elements by toll-like receptors. (23) Subsequently, the transcription factors MyD88 and IRF-7 stimulate the production of *ifn*φ, thereby activating ISGs and the production of antiviral factors, such as MxA, ISG-15, and RSAD2 (also known as viperin) in neighbouring cells. One essential transducer element in the human antiviral response is STAT1, which is part of the JAK/STAT signalling pathway. In zebrafish, two paralogues are present, *stat1a* and *stat1b*, of which *stat1a* was shown to share an ancestral virus-related cytokine-signalling mechanism similar to humans. (24, 25) The zebrafish *stat1a* and *stat1b* genes also showed differences in expression patterns, tissues specificity, and functionality. *stat1a* has a broad expression pattern at low levels throughout the whole larvae, which might indicate a virus-related function. In contrast, *stat1b* is expressed early and abundantly in the hematopoietic tissue of zebrafish embryos and plays a key role in zebrafish haematopoiesis. (25)

This study aimed to explore the zebrafish model to study RVFV replication, dissemination, and host innate immune response. We demonstrate that RVFV is able to replicate to high titers in zebrafish larvae and triggered a virus-specific innate immune response. Moreover, we reveal the virus tissue tropism in a whole-body organism for the first time, which largely recapitulates what is found in the human host and can be inhibited by a specific antiviral. Additionally, *stat1a* and *stat1b* knock-out zebrafish were created to investigate the role of the JAK/STAT pathway in RVFV infection and pathogenesis.

## Materials and methods

### Ethics statement

All animal experiments were approved and performed according to the rules and regulations of the Ethical Committee of KU Leuven (P164/2020), in compliance with the regulations of the European Union concerning the welfare of laboratory animals as declared in Directive 2010/63/EU.

### Zebrafish husbandry and transgenic lines

Routine zebrafish (*Danio rerio*) maintenance was performed under standard aquaculture conditions at 28.5 °C and a 14-hour light / 10-hour dark cycle. Fertilized eggs were collected via natural spawning. Embryos were maintained in Danieau’s solution (1.5 mM HEPES, 17.4 mM NaCl, 0.21 mM KCl, 0.12 mM MgSO_4_, and 0.18 mM Ca(NO_3_)_2_, and 0.6 μM methylene blue) under the same conditions as adults. Wildtype AB zebrafish were obtained from ZIRC (Eugene, USA). *Tg(fabp10a:DsRed; nacre*) was created as part of earlier studies in the laboratory of Peter de Witte. The *stat1a*^−/−^ and *stat1b*^−/−^ transgenic lines were generated as described below.

### Cells and viruses

Vero E6 cells (ATCC^®^ CRL-1586^™^) were cultured in Dulbecco’s Modified Eagle Medium (DMEM) (Gibco) supplemented with 10% fetal bovine serum (FBS) (Gibco), 0.075 g/L sodium bicarbonate (Gibco), and 1% penicillin-streptomycin (10 000 U/mL, Gibco). End-point titrations were performed using medium containing 2% FBS. Prof. J. Kortekaas at Wageningen University & Research kindly provided the RVFV_35/74_ strain and its reporter clone RVFV_eGFP_, which expresses eGFP from the S-segment by replacing the NSs protein locus. (26) Virus stocks were expanded using Vero E6 cells after which the titre (as CCID_50_) of the new grown stocks was determined by endpoint titration on Vero E6 cells. This, and all subsequent experiments using live virus were performed in a BSL-3 facility.

### Generation of the *stat1a* and *stat1b* CRISPR knockout zebrafish lines

The *stat1a*^−/−^ and *stat1b*^−/−^ knockout lines were generated using CRISPR/Cas9 technique following a modified protocol of Vejnar *et al.*. (27) Single guide RNAs (sgRNAs) were designed using the online tool CRISPRscan (28) targeting exon 8 of *stat1a* gene (ENSDART00000005720.6; 5’-GGCCGAGGTGTTGAACCTGG-3’) and exon 8 of *stat1b* gene (ENSDART00000000280.11; 5’-GGTCTCCAGGTTCACGGTGG-3’). The sgRNA DNA templates were synthesized via fill-in PCR using KAPA HiFi DNA Polymerase (Roche) and sgRNA-specific and universal primers (Table 1) and subsequently purified with the QIAquick PCR Purification Kit (Qiagen). sgRNAs mRNA was transcribed using AmpliScribe-T7 Flash Transcription kit (Epicentre), DNase treated, and precipitated with sodium acetate/ethanol. To optimize the Cas9 expression and its nuclear targeting in zebrafish, we used a zebrafish codon-optimized version of *S. pyogenes* Cas9 (pT3TSnCas9n, Addgene 46757). The plasmid was linearized with XbaI restriction enzyme and Cas9 *in vitro* transcription was performed using mMESSAGE mMACHINE^™^ T3 Transcription Kit (Invitrogen). IVT sgRNAs were treated with DNase and precipitated with lithium chloride/ethanol.

**Table 1:**
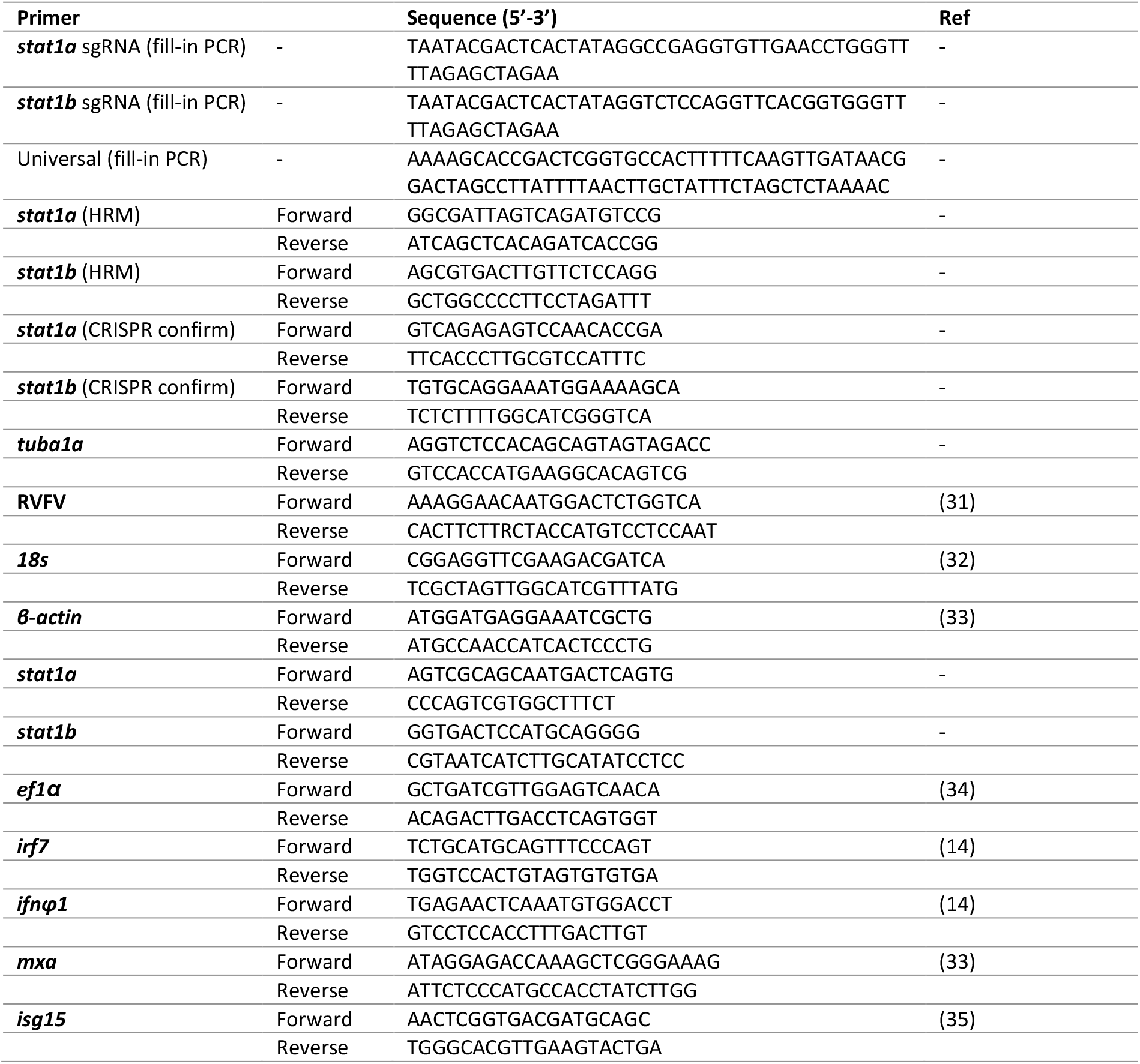
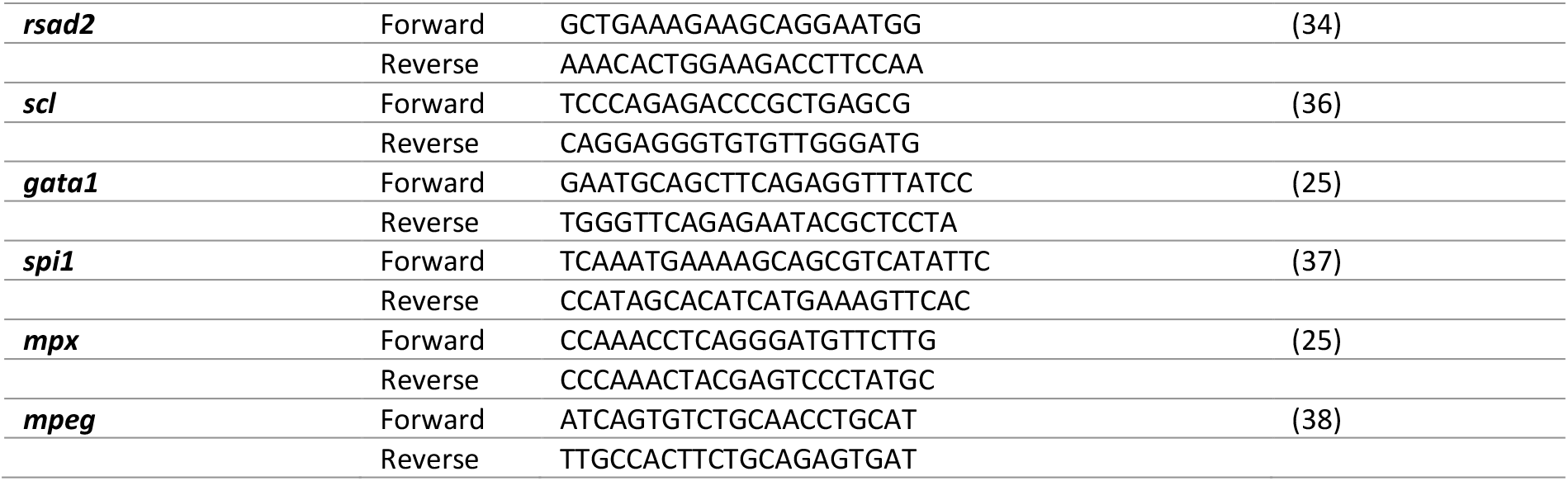
List of primers

Single cell-stage fertilized wildtype embryos of AB line were injected with a mix of 150 pg Cas9 mRNA and 100 pg *stat1a* or *stat1b* sgRNA and (in 1 nL volume) using a Femtojet 4i pressure microinjector (Eppendorf) and a M3301R Manual Micromanipulator (WPI). After the initial injections, the efficiency of CRISPR knockout was verified using high resolution melting (HRM) analysis with gene-specific primers. The remaining sgRNA/Cas9-injected embryos were raised until adulthood, outcrossed with WT adults of nacre background, and screened for indels by Sanger sequencing. F0 founders for *stat1a* and *stat1b* KO lines with germline transmission and high rate of indel mutations were selected to establish the knockout line. F1 generation embryos of F0 founders were raised to adulthood, fin clipped, and Sanger sequenced using gene-specific primers. Individuals carrying heterozygous and homozygous 10-nucleotide deletion and 8-nucleotide insertion in *stat1a* and *stat1b* gene, respectively, were identified. All experiments were performed on the F2 embryos/ larvae coming from homozygous knockout incrossing.

### RNA extraction and RT-qPCR analysis for CRISPR confirmation

Total RNA was extracted using TRIzol reagent (Invitrogen), followed by phenol-chloroform extraction, isopropanol precipitation, and ethanol washes. The resulting RNA pellet was air-dried, dissolved in nuclease-free water (Thermo Scientific), and subsequently treated with RNase-free DNase (Roche). Then, 1 μg of total RNA was reverse transcribed with the High-Capacity cDNA Reverse Transcription kit (Applied Biosystems) according to the manufacturer’s protocol. Next, the generated cDNA was diluted (1:20) and amplified using *stat1a-* or *stat1b*-specific primers and 2x SsoAdvanced Universal SYBR Green Supermix (Bio-Rad) in Hardshell^®^ Low Profile Thin-wall 96-well skirted PCR plates (Bio-Rad) on a CFX96 touch RT-PCR detection system (Bio-Rad) under cycling conditions according to the manufacturer’s protocol. The relative expression levels were quantified using the comparative Ct method (ΔΔCt) with CFX Maestro software (Bio-Rad). (29) Transcripts were normalized against the *tuba1a* reference gene using specific primers.

### High resolution melting (HRM) analysis for genotyping

To extract genomic DNA, a fin clip of a zebrafish was placed in 20 μL of 50 mM NaOH and heated at 95°C for 10 minutes, followed by neutralization using 100 mM Tris HCl (pH 8.0) (1/10 volume). Lysed samples were genotyped by performing a PCR reaction with Precision Melt Supermix for HRM analysis (Bio-Rad #172-5112) and *stat1a-* or *stot1b*-specific primers (Table 1) in a CFX96 touch RT-PCR detection system (Bio-Rad) using Hardshell^®^ Low Profile Thin-wall 96-well skirted PCR plates (Bio-Rad). Curves were analysed using the Precision Melt Analysis^™^ Software (Bio-Rad). Genotypes/ sequences of the individual larvae clustering together were confirmed with Sanger sequencing and were analysed using SeqMan software (DNASTAR Lasergene).

### RVFV injection in zebrafish larvae

At 3 dpf zebrafish larvae were anaesthetized by immersion in 0.4 mg/mL tricaine in Danieau’s solution. While sedated, the larvae were infected with RVFV_35/74_ or RVFV_eGFP_ in 1x PBS (in 2 nL) through microinjection into the pericardial cavity. Inoculated larvae were kept in Danieau’s solution under standard aquaculture conditions with up to 15 larvae per well of a 6-well plate up to 6 days post infection (dpi) and daily inspected under a stereomicroscope for clinical signs of infection (e.g., posture, development, swimming behaviour, or signs of oedema).

### Antiviral treatment

2’-Fluoro-2’-deoxycytidine (2’-FdC) (Biosynth-Carbosynth) was dissolved in 100% DMSO (VWR Chemicals). The maximum tolerated concentration (MTC) of 2’-FdC in the zebrafish larvae was determined following the protocol described by Van Dyck et al. and non-toxic concentrations were selected for antiviral testing. (30) Antiviral treatment of the RVFV-injected larvae occurred immediately after infection through immersion in 2’-FdC solution, with final concentrations of 1000 μM and 10 mM.

### Tissue homogenization and viral end-point titrations

Every dpi, ten zebrafish larvae per sample were collected in Precellys^®^ tubes and euthanized using an overdose of tricaine. The larvae were homogenized in 300 μL DMEM by homogenizer using 1 cycle of 5 sec at 6300 rpm (Precellys 24, Bertin Technologies) and centrifuged (9 000 rcf, 5 min) to pellet any debris. Infectious RVFV particles in the supernatants were quantified by end-point titrations on confluent Vero E6 cells in medium containing 2% FBS in 96-well plates. Viral titres were calculated with the Reed and Muench method using the Lindenbach calculator and were expressed as TCID50 per larvae. (31)

### RNA extraction and viral RNA quantification

Every dpi, ten zebrafish larvae per sample were collected in Precellys^®^ tubes and euthanized using an overdose of tricaine. The larvae were homogenized in 350 μL RLT-lysis buffer (Qiagen) by homogenizer using 1 cycle of 5 sec at 6300 rpm (Precellys 24, Bertin Technologies). Total RNA was extracted using the RNeasy^®^ Mini kit (Qiagen), according to the manufacturer’s protocol.

Viral RNA was detected by RT-qPCR iTaq^™^ Universal SYBR^®^ Green One-Step Kit using RVFV-specific primes (Table 1). Cycling conditions: reverse transcription at 50°C for 10 min, initial denaturation at 95°C for 3 min, followed by 40 cycles of amplification (95°C for 15 sec, 60°C for 30 sec) (Roche LightCycler 96, Roche Diagnostics). For absolute quantification, standard curves were generated using 10-fold dilutions of DNA template of known concentrations.

### Quantification of expression of innate immune-related gene expression

cDNA was generated using the ImProm-II Reverse Transcription System (Promega) by adding 1 μL of random hexamers to a total of 1 μg of extracted total RNA and incubated at 70°C for 5 min, followed by 5 min at 4°C. A mix of 8 μL of Improm II 5x reaction buffer, 6 mM MgCl_2_, 0.5 mM deoxynucleoside triphosphate, 40 units of RNase inhibitor, and 1 μL of Improm II reverse transcriptase was mixed and incubated at 25°C for 5 min, 37°C for 1 hour, and 72°C for 15 min. qPCR was performed using SsoAdvanced^™^ Universal SYBR^®^ Green Supermix and 500 nM of gene-specific forward and reverse primers (Table 1) with cycling conditions of 95°C for 3 min followed by 40 cycles of 95°C for 15 sec, 55°C for 30 sec, and 72°C for 30 sec (Roche LightCycler 96, Roche Diagnostics). Data was normalized against the housekeeping genes (*18s, β-actin*, and *ef1α*) and compared with PBS-injected zebrafish larvae to determine the fold induction of the expression, according to the 2^ΔΔCt^ method.(29)

### Whole mount immunochemistry and imaging

Zebrafish larvae from 0 to 6 dpi were anesthetized in 0.4 mg/mL tricaine in Danieau’s solution and fixed in 3.8% formaldehyde in 1x PBS. After fixation, three wash steps were performed using 1x PBS supplemented with 0.1% Tween-20 (1x PBST). From this point, the larvae were exported out of the BSL-3 facility for further processing and analyses in laboratories of lower BSL. Permeabilization was achieved starting with a wash step of 30 min in distilled H2O, followed by 20 min incubation in 100% acetone at −18°C, after which the larvae were washed three times with HBSS (Gibco), followed by immersion in collagenase (1 mg/mL in HBSS) for 60 min. After three washing steps using 1x PBST and 1% DMSO (1x PBSDT) the larvae were blocked for 1 hour in a blocking solution (10% sheep serum in 1x PBSDT) and incubated overnight at 4°C in the blocking solution containing primary antibody (Table 2). Next, the larvae were washed (4x with 1x PBSDT), with the final washing step of at least 2 hours at room temperature. An additional blocking step was done for 1 hour in the blocking solution and afterwards the larvae were incubated overnight at 4°C in the blocking solution containing secondary antibody (Table 2). The next day, the larvae were washed (3×, 1× PBST) and incubated for 30 minutes in 2 μg/mL Hoechst 33342 (Invitrogen) in 1× PBT at room temperature. After the incubation, larvae were washed six times in 1× PBST over the course of 3 hours. Images were acquired with a Leica DMI8 microscope, using 10x dry objective (NA 0.32) or 20× dry objective (NA 0.40). Image processing was done in LAS X software, version 3.7.4 (i.e., panel merging, optimal projections, and 3D deconvolution, using default set parameters).

**Table 2:** Antibodies used in whole mount immunohistochemistry

| Antigen | Host | Dilution | Source |
| --- | --- | --- | --- |
| GFP | Rabbit | 1:1000 | Chromotec (AB_2749857) |
| RFP | Mouse | 1:1000 | Chromotec (AB_2631395) |
| Anti-Rabbit 488 | Goat | 1:500 | Invitrogen (A-11008) |
| Anti-Mouse 594 | Goat | 1:500 | Invitrogen (A-11032) |

### Statistical analysis

GraphPad Prism (GraphPad Software, Inc.) was used to perform statistical analysis. To evaluate differences between means, two-tailed unpaired Student *t*-test was used. To determine normal distributions, Shapiro-Wilk tests were performed. Nonparametric data was evaluated using the Kruskal-Wallis test. Survival studies were analysed using Log-rank (Mantel-Cox) test. Values *P*<0.05 are considered statistically significant (\**P*<0.05; \*\**P*<0.01; \*\*\**P*<0.001; \*\*\*\**P*<0.0001; ns, not significant).

## Results

### RVFV replicates in zebrafish larvae and induces an innate immune response

To establish a zebrafish larvae infection model for RVFV, we injected ^~^1 CCID_50_ of RVFV_35/74_ (in a 2 nL volume) or a UV-irradiated sample of the same virus in the pericardial cavity of 3 dpf larvae (Figure 1A). This site of injection was selected as it mimics the human transmission route through a mosquito bite. RVFV_35/74_ viral RNA was detected from 1 dpi onwards, reaching a peak of ^~^4 log_10_ RNA copies per larva at 3 dpi (Figure 1B). In the following days, viral RNA load slowly decreased to ^~^2 log_10_ RNA copies per larva at 6 dpi. When the larvae were injected with the UV-irradiated inoculum, no increase in viral RNA, hence no viral replication was observed (Figure 1B). Additionally, infectious virus titres were determined by endpoint titration to confirm the presence of infectious virus progeny. The infectious virus titres coincided with the detected RNA levels, with a peak observed at 3 dpi, confirming that viral RNA detection correlates with the presence of infectious virions (Figure 1C). No signs of disease (emaciation, skeletal deformation, oedema, or haemorrhages) were detected in the RVFV-injected larvae at any time during the experiment and no difference in mortality was observed between the larvae inoculated with virus or PBS as control (S. Figure 1A).

**Figure 1.**
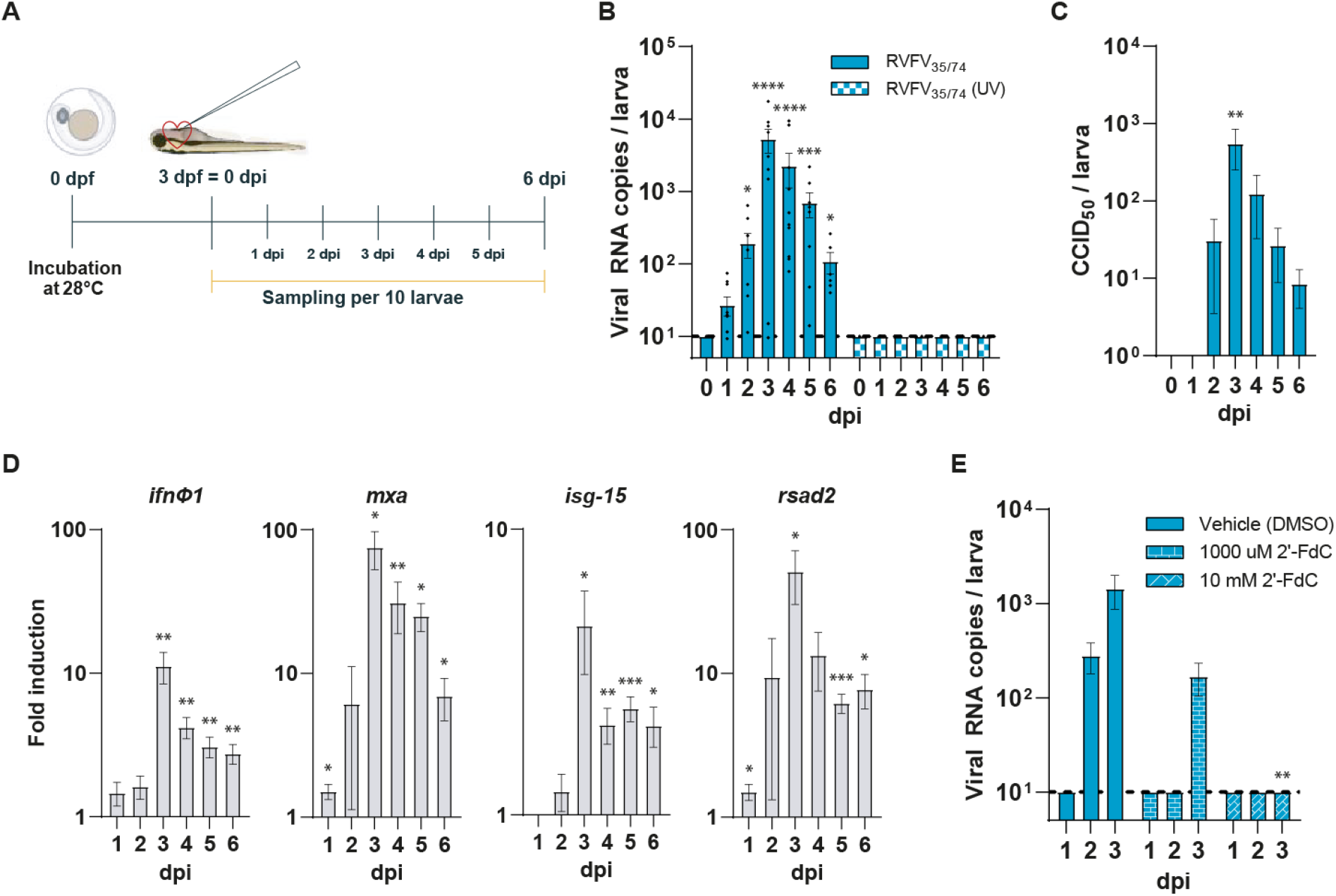
RVFV infection in zebrafish larvae resulting in viral replication and innate immune response. **A)** Schematic representation of the experimental timeline, conditions, and injection site. Drawings acquired through BioRender. **B)** Viral RNA copies of RVFV_35/74_ per zebrafish larva. Mean ± s.e.m. of 5-8 pools of 10 larvae from 8 independent experiments. The dotted line shows LOQ. **C)** Titres of RVFV_35/74_ infectious viral particles in zebrafish larvae. Mean ± s.e.m. of 6 pools of 10 larvae from 6 independent experiments. **D)** Relative fold induction of *ínfφ1*, *mxa*, *isg-15*, and *rsad2* in RVFV_35/74_ infected zebrafish larvae compared to PBS-injected zebrafish larvae. Results determined by RT-qPCR and normalized to *18s*, *ef1α*, and *β-actin* at 0 dpi. Mean ± s.e.m. of 8 pools of 10 larvae from 8 independent experiments. **E)** Viral RNA copies of RVFV_35/74_ per zebrafish larva after immersion in Danieau’s solution containing the vehicle, 1000 μM, or 10 mM 2’-FdC at 0 dpi. Mean ± s.e.m. of 5 pools of 10 larvae from 5 independent experiments. The dotted line shows LOQ. dpf = days post fertilization; dpi = days post infection; RVFV = Rift Valley fever virus; 2’-FdC = 2’-Fluoro-2’-deoxycytidine; *P<0.05; **P<0.01; ***P<0.001; ****P<0.0001; ns = not significant

To demonstrate that zebrafish mount an innate immune response upon infection with RVFV, we quantified the mRNA expression levels of innate immunity-related genes. Upon infection with RVFV_35/74_, the expression of interferon-φ1 (*ifnφ1*) and downstream IFN-stimulated genes *mxa*, *isg-15*, and *rsad2* were increased with maximum levels coinciding with the peak of virus replication at 3 dpi (Figure 1D).

To validate our model for antiviral studies and generate additional evidence that RVFV can effectively replicate in zebrafish, we treated RVFV_35/74_ infected larvae with 2’-Fluoro-2’-deoxycytidine (2’-FdC), a nucleoside polymerase inhibitor with a proven *in vitro* and *in vivo* antiviral effect against RVFV. (39) Immediately after virus infection (0 dpi), the larvae were exposed to 10 mM 2’-FdC, after which no viral replication was detected for the first 3 dpi. When a 10-fold lower concentration of 2’-FdC was used, no viral RNA was detected at 1 and 2 dpi, while at 3 dpi viral RNA levels were detectable but still lower than those found in DMSO-treated controls (Figure 1E).

Taken together, we demonstrated that zebrafish larvae are permissive to RVFV infection and viral replication, which induced a virus-induced innate immune response. Moreover, treatment with an antiviral compound completely prevented RVFV replication, demonstrating the utility of the zebrafish model for testing and identification of novel small molecule inhibitors of RVFV replication.

### RVFV has a tropism for the liver and sensory nervous system in zebrafish larvae

To investigate the tropism of RVFV upon infection in zebrafish larvae, we used an eGFP fluorescently labelled RVFV_35/74_ strain, hereafter named RVFV_eGFP_. Infection with ^~^15 CCID_50_ of RVFV_eGFP_ in the pericardial cavity of the larvae yielded similar replication kinetics, but with an earlier peak of replication of ^~^3 log_10_ RNA copies per larva at 2 dpi (Figure 2A). Similarly, as in experiments with the non-labelled virus, there was no difference in mortality between RVFV_eGFP_ infected larvae and PBS-injected controls throughout the experiments (Figure S1A). By means of whole mount immunohistochemistry and fluorescence microscopy, we were able to image RVFV_eGFP_ infected zebrafish larvae starting from 1 dpi (Figure 2B). After infection, most larvae showed RVFV_eGFP_ positive signal around the site of injection at 1 or 2 dpi and developed an infection of the liver in a spot-like pattern during early days of the infection that became more widespread at 3-6 dpi. The virus accumulation in the liver was further investigated by RVFV_eGFP_ infection of a transgenic zebrafish line, *Tg*(*fabp10a:DsRed; nacre*), expressing DsRed fluorophore in the hepatocytes. Upon infection, we observed an overlap of the red fluorescent signal from the hepatocytes and the eGFP translated from the viral genome (Figure 2C), confirming that the liver is the main target organ for RVFV replication in zebrafish, similar to humans. In two out of six independent experiments, nearly all larvae showed infection of their sensory nervous system, including the optic tectum, the anterior and posterior macula, the lateral line including the related ganglia, and the retina (Figure 2D). The typical pattern of the neuromasts belonging to the lateral line around the head and along the body of the larvae was clearly distinguishable. Out of 30 RVFV-positive larvae we found two larvae that did show infection of the optical tectum and the most proximal neuromasts, but without having spread to the more distal parts of the lateral line (Figure S2). Remarkably, none of the larvae showed an overlapping infection pattern including both the liver and the sensory nervous system.

**Figure 2.**
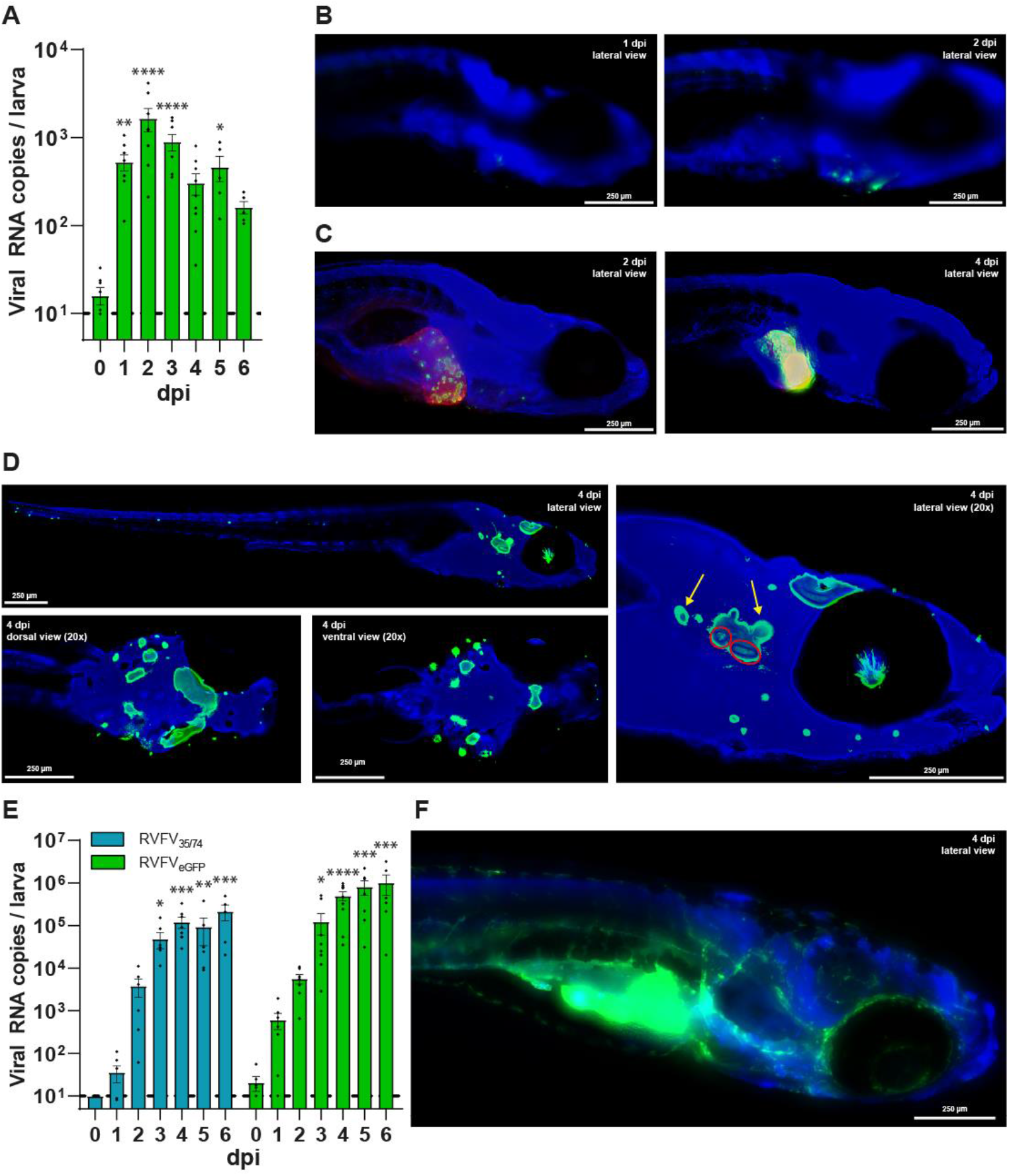
Viral replication and dissemination of RVFV_eGFP_ in zebrafish larvae, with infection of the liver, sensory nervous system and vascular system. **A)** Viral RNA copies of RVFV_eGFP_ per zebrafish larva. Mean ± s.e.m. of 6-8 pools of 10 larvae from 8 independent experiments. **B-D, F)** Whole mount immunohistochemistry fluorescence images of RVFV_eGFP_ infected zebrafish larvae at 10x magnification, using anti-GFP primary antibody and Hoechst 33342. **B)** Wildtype zebrafish larva at 1 and 2 dpi showing progressing RVFV infection around the site of injection. **C)** *Tg*(*fabp10a:DsRed; nacre*) zebrafish larvae demonstrating infection in the liver using anti-RFP primary antibody. **D)** Wildtype zebrafish larva showing infection of the retina, optic tectum, the anterior and posterior macula (red circle), the ganglia (yellow arrows), and neuromasts of the lateral line. dpi = days post infection **E)** Viral RNA copies of RVFV_35/74_ and RVFV_eGFP_ per zebrafish larva immersed in Danieau’s solution supplemented with ruxolitinib (25 μM) from 0 dpi. Mean ± s.e.m. of 8 pools of 10 larvae from 8 independent experiments. **F)** Wildtype zebrafish larva at 6 dpi, showing extensive dissemination of RVFV_eGFP_ infection after immersion in Danieau’s solution supplemented with ruxolitinib (25 μM) from 0 dpi. Image has not been 3D-deconvoluted. dpi = days post infection; RVFV = Rift Valley fever virus; *P<0.05; **P<0.01; ***P<0.001; ****P<0.0001; ns = not significant

To confirm the role of the JAK/STAT signalling pathway in RVFV infection we exacerbated the viral infection by introducing the JAK signalling inhibitor ruxolitinib to the swimming water of the zebrafish larvae. Viral RNA levels were increased upon infection with both the RVFV_35/74_ strain and the RVFV_eGFP_ reaching ^~^5 log_10_ RNA copies/larva at 3 dpi (Figure 2E). Beyond that timepoint viral RNA levels continued to increase to values close to ^~^6 log_10_ RNA copies/larva at 6 dpi, which contrasts with the ^~^2 log_10_ RNA copies/larva found in the above shown regular infection. This observation supports a key role for the JAK/STAT pathway in the host response to the infection in zebrafish larvae. Imaging of RVFV_eGFP_ infected larvae at 6 dpi showed extensive dissemination of the virus. Their abdominal area was bright green as the virus had infected the liver, large portions of the abdominal cavity and gastrointestinal tract, and their entire vascular system (Figure 2F). While eGFP signal was detected throughout the head and the area surrounding the eye, the pattern of observed fluorescence indicates that it was not the neurons that were infected, but rather the vascular system. In fact, none of the larvae showed any signs of infection in the sensory nervous system, lateral line, or retina. Moreover, despite the more extensive replication and dissemination of the virus in larvae treated with the JAK inhibitor, their survival rates were not affected (Figure S1B).

### Inhibiting the JAK/STAT pathway exacerbates RVFV replication and dissemination in zebrafish larvae

As *stat1* is an important signal transducer in the JAK/STAT signalling pathway, its loss of function was investigated in zebrafish. Due to genome duplication in zebrafish, two paralogues of the human *STAT1* gene exist i.e., *stat1a* and *stat1b* and thus two zebrafish knockouts were generated. Using CRISPR/Cas9 genome editing technology (40), we targeted the 8^th^ exon of the zebrafish *stat1a* and *stat1b* genes encoding a part of the ‘coiled-coil’ domain of the respective proteins. For the *stat1a* gene, a positive founder transmitting 10-nucleotide frameshifting deletion was selected and for the *stat1b* gene we chose a positive founder carrying 8-nucleotide insertion, both would result in a premature stop codon and consequently truncated proteins (Figure 3A). The genomic sequence of both *stat1a* and *stat1b* mutant larvae was confirmed by Sanger sequencing and the selected founders were crossed to obtain the homozygous larvae in the F2 generation. These F2 homozygotes survived until adulthood, were fertile, and the F3 offspring from the homozygous parents was used for all subsequent experiments. Quantification of mRNA expression showed a significant reduction in homozygous larvae (Figure 3B & 3D), likely resulting from nonsense-mediated mRNA decay. In the *stat1b*^−/−^ larvae, a decrease in *stat1b*^−/−^ mRNA expression was confirmed and accompanied by a remarkable increase of *stat1a* expression of ^~^1.5-fold. No obvious morphological defects were observed in these F3 generation *stat1a*^−/−^ and *stat1b*^−/−^ larvae. Survival studies during raising to adulthood demonstrated that *stat1a*^−/−^ larvae had a decreased survival rate, likely due to a higher susceptibility to infections while the survival of *stat1b*^−/−^ larvae was not affected (Figure 3C & 3E).

**Figure 3.**
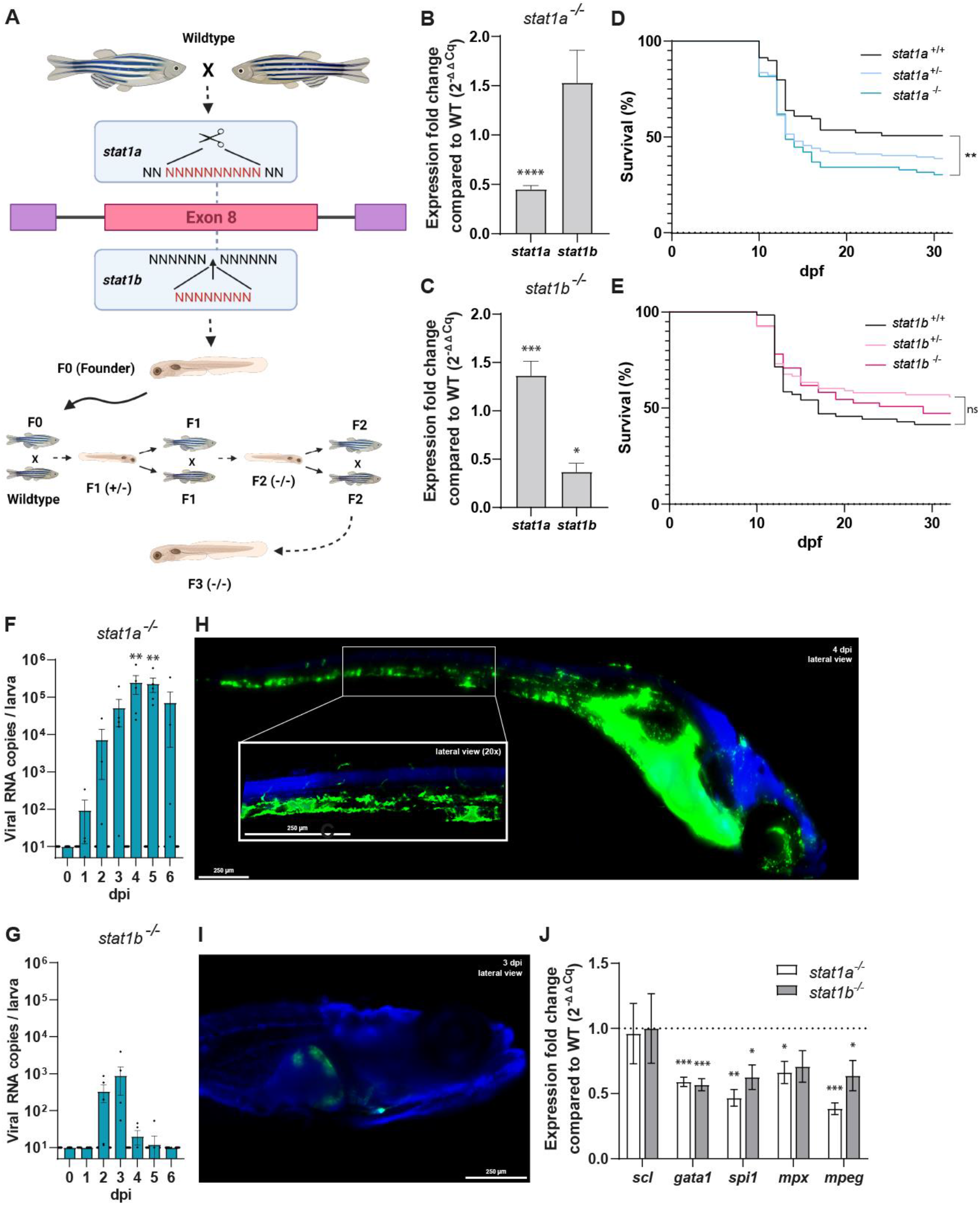
The effect of *stat1a* and *stat1b* knockout on RVFV viral replication, dissemination, and innate immune response. **A)** Schematic representation of the creation of *stat1a*^−/−^ and *stat1b*^−/−^ larvae. Created in BioRender. **B)** The relative mRNA expression fold change of *stat1a* and *stat1b* in *stat1a*^−/−^ zebrafish larvae. **C)** The relative mRNA expression fold change of *stat1a* and *stat1b* in *stat1b*^−/−^ zebrafish larvae. **D)** Survival curves of *stat1a^+/+^* (n=69)*, stat1a^+/-^* (n=129)*, and stat1a*^−/−^ (n=74) larvae of the F2 generation. **E)** Survival curves of *stat1b^+/+^* (n=71)*, stat1b^+/−^* (n=92), and *stat1b*^−/−^ (n=55) larvae of the F2 generation. **F)** Viral RNA copies of RVFV_35/74_ per *stat1a*^−/−^ zebrafish larva. Mean ± s.e.m. of 5-6 pools of 10 larvae from 6 independent experiments. **G)** Viral RNA copies of RVFV_35/74_ per *stat1b*^−/−^ zebrafish larva. Mean ± s.e.m. of 6 pools of 10 larvae from 6 independent experiments. **H,I)** Whole mount immunohistochemistry fluorescence images of RVFV_eGFP_ infected *stat1a^−/−^* (H) and *stat1b*^−/−^ (I) zebrafish larvae at 10x magnification, using anti-GFP primary antibody and Hoechst 33342. *Stat1a^−/−^*zebrafish larva at 6 dpi, showing extensive dissemination of RVFV_eGFP_ infection. *Stat1b*^−/−^ zebrafish larva at 6 dpi, showing limited dissemination of RVFV_eGFP_ infection. **J)** Fold induction of gene expression of hematopoietic markers in *stat1a*^−/−^ and *stat1b*^−/−^ larvae relative to wildtype larvae at 6 dpf. Results determined by RT-qPCR and normalized to *18s, ef1α*, and *β-actin* mRNA. dpi = days post infection; RVFV = Rift Valley fever virus; WT = wildtype; *P<0.05; **P<0.01; ***P<0001; ****P<0.0001; ns = not significant.

Upon injection of the *stat1a*^−/−^ larvae with RVFV_35/74_, we observed a similar course of infection as for wildtype larvae treated with ruxolitinib, with viral loads above 5 log_10_ RNA copies per larva (Figure 3F). Similarly, viral RNA levels did not rapidly diminish after 3 dpi but remained stable until the end of the experiment. In addition, the dissemination pattern of RVFV_eGFP_ seems analogous to ruxolitinib treated larvae, with very bright signal in the liver and widespread infection of the vascular system (Figure 3G). While virus dissemination is extensive, there was no effect on the survival rates during the course of the experiment (Figure S1C). Replication of RVFV_35/74_ was impaired in *stat1b*^−/−^ larvae with only minor levels of viral RNA detected at 2, 3 and 4 dpi (Figure 3H). Regarding the dissemination of RVFV_eGFP_, no changes in site of replication were detected; the eGFP signal was mainly found around the place of injection and in few cases in the liver (Figure 3I). The overexpression of *stat1a* in *stat1b*^−/−^ larvae could explain, at least in part, the lowered RVFV replication. Besides, *stat1b* was described to play a role in haematopoiesis, promoting myeloid development at the expense of erythroid development. (25) We hypothesized that the absence of *stat1b* conditioned the number of monocyte/macrophages present in the larvae, which affected the ability of the virus to infect and propagate within the larvae. In fact, it was confirmed that cells of the myeloid lineage are a host for RVFV and a determining factor in viral dissemination. (41) Hence, we investigated potential differences in the leukocyte population in both *stat1a*^−/−^ and *stat1b*^−/−^ larvae compared to wildtype larvae at 6 dpf. We used *scl* as a marker for hematopoietic progenitor cells, *gata1* for erythrocytes, *spi1* (*pu1*) for myeloid precursors, *mpx* for heterophil granulocytes, and *mpeg* for monocytes. In line with what was suggested by Song *et al*., we detected low levels of the myeloid lineage precursor marker *spi* and both *mpx* and *mpeg* markers (Figure 3J). However, these levels were the same in the *stat1a*^−/−^ line, where *stat1b* is abundantly present. Moreover, we detect lower levels of the erythrocyte marker *gata1* in both knockout lines compared to wildtype, which is also not in line with the expected shift of haematopoiesis towards erythropoiesis. Only the marker for hematopoietic progenitor cells the *stat1a*^−/−^ and *stat1b*^−/−^ larvae showed similar levels compared to wildtype larvae.

## Discussion

Here, we report a new infection model for RVFV using zebrafish larvae. In this model we observed extensive viral replication and dissemination to organs compatible with what has been described in the human host. While mouse models for RVFV have already been described, zebrafish as a small vertebrate model is a worthwhile endeavour and will aid in unravelling the remaining knowledge gaps in the field. Even though zebrafish are evolutionary more distant to humans than rodents, they have high genetic similarity with approximately 82% of human disease-related genes having at least one orthologue in zebrafish. (42) On top of the higher compliance with the ethical 3Rs rule (particularly when used before 120 hours post fertilization), zebrafish are specifically advantageous for high biosafety level (BSL) environments due to their small size, ease of manipulation and transparency. Their small size enables the use of 96-well plates for housing and thereby generates lower volumes of biohazard infectious waste compared to rodents (cages, bedding, food, water, cadavers, etc.). Moreover, virus injection procedures do not require the use of sharps, eliminating the risk of needle stick injuries, which is particularly relevant in this high BSL setting. Additionally, zebrafish transparency facilitates the visualisation of viral dissemination throughout the course of infection with a reporter virus and thus provides novel insights into RVFV infection and disease.

Over the years, zebrafish have been used for the identification of bioactive compounds, toxicity studies and beyond, both in manual and automated settings. (43, 44) Here we studied the *in vivo* efficacy of the known broad-spectrum viral polymerase inhibitor, 2’-FdC, active against RVFV infection in mice. (39) The treatment proved to be efficient, yielding ^~^2 log_10_ reduction in virus progeny at the highest tested concentration. In this way, we showed that antiviral activity was also recapitulated in a zebrafish larval model and thus can be used for discovery of other (novel) antiviral drugs. Other advantages of the zebrafish model include the capacity to immediately and easily monitor compound toxicity in a complete model organism, and the low amounts of compound needed for drug testing. (44) Moreover, if the solubility of the compound allows, it can be simply added to the swimming water without specific formulation. This demonstrates that zebrafish models are simple and affordable tool organisms, which can be used early in a drug discovery campaign to generate invaluable high-quality *in vivo* data by using very limited resources.

Upon injection of RVFV into 3 dpf zebrafish larvae, the virus was able to replicate efficiently to high titres. Pericardial injection was chosen as the way of inoculation to mimic the mosquito bite route of infection by directly releasing the virus into the bloodstream of the larva. The infection progressed rapidly with a swift rise in viral titres, akin to what is observed in studies using mice. (7) After the infection peaked at 3 dpi, we observed a gradual decrease in viral RNA and infectious particles, which can be attributed to the innate immune response mounted by the larvae. IFN plays an important role in the immunity against infections by negative-strand RNA viruses, including RVFV. (45) The involvement and close relationship between the IFN-competency of the host and its immunogenic response to RVFV infection is well established. The ability of RVFV to inhibit IFN pathway functionality correlates with increased viremia. (46, 47)

We also demonstrated that in zebrafish larvae *ifnφ* and several downstream ISGs are strongly induced early upon infection, successfully controlling the RVFV infection. The contribution of the innate immune system can be assured, as the adaptive immunity is not functional yet at this stage in larval development. (18)

Using an RVFV_eGFP_ we were able to visualise the RVFV infection and dissemination throughout the body of the larvae. This reporter virus lacks the NSs gene, which is replaced by the *eGFP* gene that is expressed upon infection of a host cell. The lack of the NSs protein might have attenuated the virus as inoculum of comparable titres to the wildtype virus did not consistently provoke an infection in the larvae. For this reason, we resorted to a 10-fold higher inoculum in all further experiments. We believe this increase in inoculum was the reason that the virus was able to reach substantial RNA levels a day earlier, when compared to our experiments using wildtype virus. We found that small spots of eGFP signal could be detected as early as 1 dpi around the site of infection in most of the larvae. The infection did not progress identically in all RVFV_eGFP_-infected larvae. At 2 dpi the majority of the larvae only showed infection around the site of injection, while in others the infection did already reach the liver. This is of particular interest since RVFV is known to be hepatotropic in both human patients and mouse models. (48) In the subsequent days, an increasing number of larvae displayed infection of the liver at varying degrees of severity. Remarkably, from 2 dpi onwards we observed infection of the sensory nervous system of the larvae from two out of six independent experiments. So far, our ability to pinpoint how and where the virus transfers from the liver or vascular system into the sensory nervous system was hampered due to the limited experimental setup in the BSL-3 environment. Live imaging could help to investigate this further in the future. Since some larvae showed infection restricted to the optic tectum, without it having spread (yet) to the lateral line or retina we speculate that the optic tectum might be the gateway for the virus to spread into the sensory nervous system and progressively to other sensory neuron-rich structures. The remaining 28 larvae displayed infection throughout their broader sensory nervous system, showing fluorescent signal in all related structures, i.e., the optic tectum, the retina, the anterior and posterior macula, the ganglia, and the neuromasts of the lateral line. These structures have been shown to be connected by ensembles of neurons within the tectum that respond to sound, flow, or visual stimuli. (49) This co-localization of neural clusters might provide the virus with an excellent site from where to disseminate and travel along afferent neuronal axons, afterwards infecting the innervated structures. Interestingly, we observed the presence of the virus in a branched structure within the eye suggesting infection and potential damage to the eye or optic nerve, which might potentially be linked to the RVFV-related blindness in humans. (50, 51)

Even though the virus is able to infect sensory nervous system-related cells and tissues, it does not seem able to make the jump into the central nervous system, since in none of our experiments we observed infection of the brain or the spinal cord. While the maturation of the zebrafish BBB occurs between 3 and 10 dpf, expression of high resistance tight junction proteins claudin-5 and ZO-1 is already present at 3 dpf, leading to reduced permeability of the brain microvessels. (52) Our finding is contradictory to earlier studies in RVFV-infected rats and ferrets, where acute encephalitis and infection of the brain was observed. (53, 54) However, it should be noted that in those studies the animals were infected through aerosol exposure, rather than via intravenous or subcutaneous administration, which alters the pathogenesis of RVFV. (55) A similar approach, i.e., RVFV inoculation through the olfactory system, could be used in zebrafish adults and eventually, in larvae. (52)

Immersion in water with the JAK inhibitor ruxolitinib exacerbated the infection and hampered its termination due to the inhibition of the JAK/STAT pathway. A very intense fluorescent signal was detected in the liver, which interfered with the imaging of the surrounding tissue. Likewise, the complete vascular system of the larvae lit up green, including microvasculature in and around the eyes, the brain, and the muscles. Despite this massive spread in the blood vessels, there was no fluorescent signal observed within surrounding neuronal, muscular, or connective tissue. Interestingly, no fluorescent signal was observed in the sensory nervous system, which supports a concept of an immune-dependent dissemination of the virus in larvae with a pharmacologically inhibited JAK/STAT pathway.

To further study the role of the JAK/STAT pathway and more specifically of its transducer STAT1 in RVFV infection, we generated two knockout zebrafish lines, *stat1a*^−/−^ and *stat1b*^−/−^. As expected, the *stat1a*^−/−^ larvae showed a similar infection progression as larvae treated with ruxolitinib. This confirms that *stat1a* is indeed a key factor in triggering the host innate immune response against RVFV infection. In *stat1a*^−/−^ larvae the virus encountered minimal resistance and was able to disseminate extensively throughout the liver and vascular system. Remarkably, despite the high viral loads, none of our experiments showed infection of any component of the sensory nervous system, either when using ruxolitinib or *stat1a*^−/−^ larvae.

When infecting *stat1b*^−/−^ larvae, much lower viral loads were detected, with the majority of larvae showing eGFP signal only at the site of infection and a small fraction showing infection of the liver. This could be explained by the significantly increased levels of *stat1a* found in these larvae, which seems to arise as compensation for the *stat1b* knockout. Such a genetic compensation in response to a gene knockout has already been described many times in several model organisms. (56) Since *stat1a* and *stat1b* genes are paralogues, they are thought to serve distinct functions, thus this apparent connection was a surprising finding. The higher levels of *stat1a* in *stat1b*^−/−^ larvae might account for the lower viral load and therefore directly lessened viral dissemination, by simply boosting the downstream-hosted innate immune response. Additionally, the reduced replication of RVFV in the *stat1b*^−/−^ larvae could be played in hand by a lower number of cells of the myeloid lineage, as indicated by low levels of *spi*, *mpx*, and *mpeg* markers. Since myeloid cells were present in reduced numbers, RVFV might have fewer opportunities to hide from the host innate immune response, which resulted in limited viral replication and more efficient infection control. While the levels of myeloid cell markers were also lower in the *stat1a*^−/−^ larvae, the virus could replicate in an unhindered manner due to the deficient host innate immune response.

Here we show that zebrafish larvae are an excellent *in vivo* model to study RVFV replication, dissemination, and host innate immune response. Their versatility and ease of use make it a well-suited tool for high biosafety environments. Our results reveal that RVFV-induced blindness likely occurs due to direct damage to the eye and peripheral neurons, rather than to the central nervous system, and that the virus does so in a *stat1* dependent manner.

This new RVFV model could be used to further dissect the interplays between virus and host and help to confirm the proposed association between innate immune gene polymorphisms and RVFV-associated clinical outcomes. (57) Also, the intricate innate immunity mechanisms activated to combat the RVFV infection that seem to directly influence which organs are affected can now be explored, thereby contributing to a better understanding of the pathogenesis of RVRV infection and to the discovery of efficacious therapies.

## Acknowledgements

We wish to thank Prof. Jeroen Kortekaas for providing us with RVFV and reporter virus suspensions, Kevin Longin, Tina Van Buyten and Jasper Rymenants for their technical assistance, Elisabeth Heylen and Sonia Mertens for their impeccable management and maintenance of the BSL-3 facility, and Kathleen Lambaerts for her care of the aquatic facility.

## Competing interests

All authors declare to have no competing interests.

## Author contributions

Conceptualization: StH, JRP

Methodology: StH, JRP, AS, PdW

Experimental work: StH, AS, ASDM, LC

Creation of zebrafish knockout lines: StH, AS, ASDM, AC

Data analysis: StH, AS, ASDM, JRP

Visualization: StH, AS

Writing: StH, AS, JRP

Review & Editing: StH, JRP, AS, ASDM, AC, LC, PdW, JN

Supervision: JRP, PdW, JN

## Funding

StH and the research leading to this work were funded by KU Leuven internal funds (C24/18/080) to JRP and PdW. ASDM was funded by Fund for O6260 Research Foundation–Flanders (FWO; 11F2919N).

## Notes

### Competing Interest Statement

The authors have declared no competing interest.

