## Supplementary material for "The dissemination of Rift Valley fever virus to the eye and sensory neurons of zebrafish larvae is *stat1* dependent": Figure S1

### Supplementary figure 1

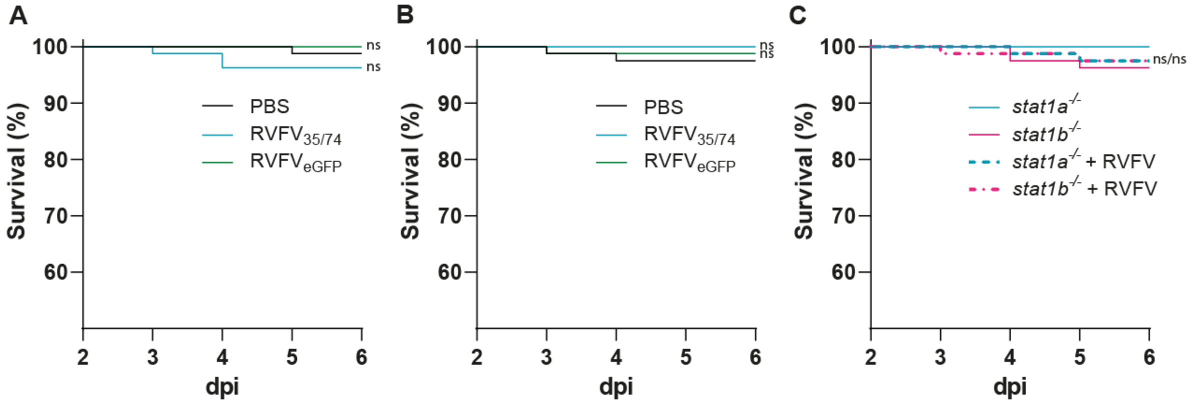


**Figure S1 - Survival curves of RVFV infected zebrafish larvae**. **A)** Survival of the larvae injected with $RVFV_{35\text{-}74}$, $RVFV_{eGFP}$ or PBS (uninfected). No significant larval death can be observed over the entire duration of the experiment. **B)** Survival of larvae infected with $RVFV_{35\text{-}74}$, $RVFV_{eGFP}$ or PBS (uninfected) while treated with ruxolitinib (continued immersion in Danieau's solution supplemented with 25 µM ruxolitinib). No significant larval death can be observed over the entire duration of the experiment. **C)** Survival of *stat1a*$^{-/-}$ and *stat1b*$^{-/-}$ larvae injected with $RVFV_{35\text{-}74}$ or PBS (uninfected). No significant larval death can be observed over the entire duration of the experiment. Statistical analysis between different groups was performed using Log-rank (Mantel-Cox) test where $p<0.05$ is considered significant. Data pooled from four independent experiments, total included numbers per group: n = 80. dpi = days post infection; RVFV = Rift Valley fever virus; ns = not significant

### Supplementary figure 2

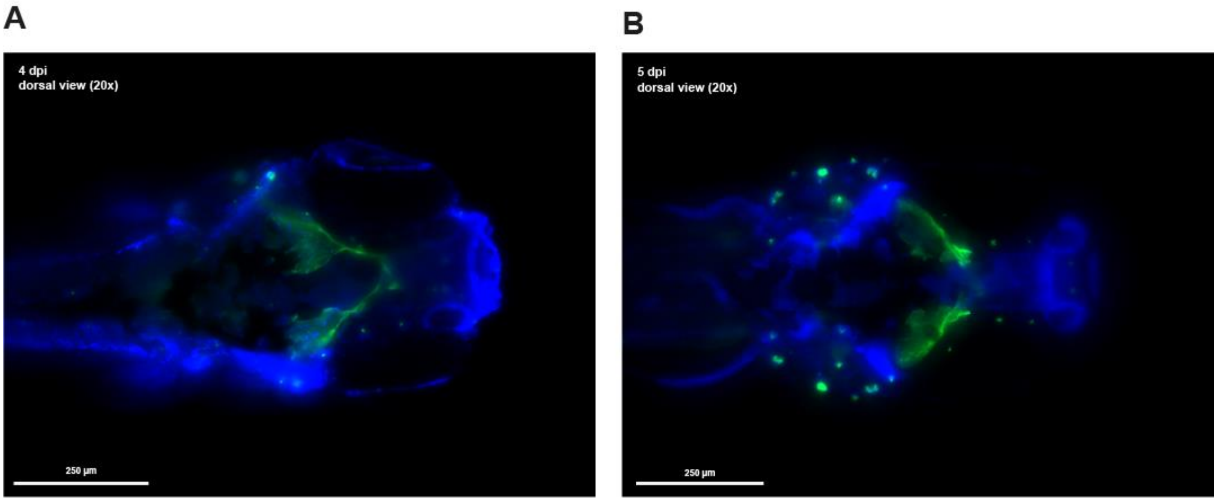


**Figure S2 – Images of $RVFV_{eGFP}$ infected larvae where the infection has spread only limited beyond the optic tectum**. Whole mount immunohistochemistry fluorescence images of $RVFV_{eGFP}$ infected zebrafish larvae at 20x magnification, using anti-GFP primary antibody and Hoechst 33342. Images were not 3D-deconvoluted. **A)** $RVFV_{eGFP}$ infected zebrafish larva at 4 dpi showing only infection of the optic tectum and the most proximal neuromasts. **B)** $RVFV_{eGFP}$ infected zebrafish larva at 5 dpi showing infection in the optic tectum and proximal neuromasts.
